# SHLD2 loss is a synthetic vulnerability to Polθ inhibition combined with radiotherapy

**DOI:** 10.1101/2025.07.04.662969

**Authors:** Gonzalo Rodriguez-Berriguete, Purusotha Thambiayah, Alessandro Cicconi, Nicole Machado, Celia Gotorbe, David Nderitu, Wei-Chen Cheng, Marie Laure Boursier, Aurora Cerutti, Vera Grinkevich, Bethany Rebekah Hill, Katjuša Koler, Sophie Alice Langdon, Jayesh B Majithiya, Suraj Menon, Shaun Moore, Joana Neves, Natalie M Palmer-Deverill, Eeson Rajendra, Marina Roy-Luzarraga, Asmita Thapa, Robert A Heald, Graeme C M Smith, Helen M R Robinson, Marco Ranzani, Geoff S Higgins

## Abstract

DNA polymerase theta (Polθ) plays a crucial role in the repair of DNA double-strand breaks (DSBs) by microhomology-mediated end joining (MMEJ). We previously demonstrated that Polθ inhibition (Polθi) is an effective and well-tolerated approach to sensitise tumours to radiotherapy (RT). Here, we profiled 54 cancer cell lines and found that Polθi induces significant radiosensitisation in most models, though with marked variability not explained by indicators of Polθ activity. To pinpoint molecular determinants of radiosensitisation by Polθi, we performed a CRISPR knockout screen which revealed loss of the TP53BP1/Shieldin pathway component *SHLD2* (*FAM35A*) as a vulnerability to Polθi combined with RT. We demonstrated that *SHLD2* loss not only increases sensitivity to RT alone, but also enhances the radiosensitising effect of Polθi, both *in vitro* and *in vivo*. Importantly, we found that *SHLD2* is deleted in a subset of human prostate cancers, often co-occurring with *PTEN* loss, an adverse prognostic factor. Furthermore, we show that *SHLD2*-deficient cancer cells are more reliant on Polθ to prevent DSB accumulation and chromosomal instability. In summary, we discovered *SHLD2* loss as a novel collateral vulnerability that can be exploited through combined treatment with Polθi and RT.

## INTRODUCTION

Radiotherapy (RT) plays a key role in the treatment of many solid tumours. When given with curative intent, RT is often combined with conventional chemotherapies to sensitise tumours to ionising radiation. However, these effects are not tumour-specific and the benefit of these treatments in improving tumour control is offset by side effects caused by increased damage to surrounding normal tissues^1^. Therefore, there is major interest in combining RT with drugs that exert a radiosensitising effect selectively in tumour cells. In this regard, we have previously reported that pharmacological inhibition of Polθ is an effective and safe approach for tumour-selective radiosensitisation in preclinical models^2^.

Polθ (encoded by the *POLQ* gene) is a type A DNA polymerase that also contains a helicase domain, and plays a critical role in the repair of DNA double-strand breaks (DSBs) by microhomology-mediated end joining (MMEJ), an error prone repair pathway^3^. During MMEJ repair, MRE11/CtIP-mediated initiation of 5′→3′ resection at the DSB generates 3′ overhangs, exposing short, complementary microhomologies^4^.

The helicase domain of Polθ then facilitates the annealing of these microhomologies, and its polymerase domain fills in and seals the resulting gaps^5,6^. This error-prone repair leaves characteristic genomic ‘scars’ at the DSB site, consisting of short deletions flanked by microhomologies, which reflect the mutagenic signature of Polθ^7^.

Polθ is synthetically lethal with components of the homologous recombination (HR) pathway, such as BRCA1 and BRCA2^8–10^. In addition, Polθi has been shown to enhance the antitumour effect of PARP inhibition (PARPi) in HR-deficient cancer cells^8,10–12^. This has led to ongoing clinical testing of Polθi in patients with HR-deficient cancers as a monotherapy and in combination with PARPi (NCT05898399, NCT04991480, NCT06077877, NCT05687110, NCT06666270, NCT06545942, NCT06560632). In contrast, we have previously demonstrated that HR deficiency is not required for Polθi-induced radiosensitisation^2^. Therefore, the combination of Polθi with RT may benefit a broader patient population than Polθi alone or in combination with PARPi. Nonetheless, it remains unclear how variability among tumours influences the radiosensitising effect of Polθi and, to date, no clinically actionable molecular markers exist to identify which patients could benefit the most from Polθi combined with RT. Addressing these gaps will be crucial for the effective clinical translation of this combination treatment.

## RESULTS

### Polθi-induced radiosensitisation varies across cancer cell lines and does not correlate with indicators of Polθ activity

Our previous work showed that small-molecule, allosteric inhibitors of Polθ’s polymerase domain –ART558 and ART899– are effective, tumour-selective radiosensitisers in preclinical models^2^. To interrogate to what extent tumour cell heterogeneity impacts radiosensitisation by Polθi and to explore its potential for broad clinical application, we evaluated ART558-mediated radiosensitisation in a panel of 54 lung, colorectal, and head and neck cancer cell lines. As shown in Figure 1A, the magnitude of radiosensitisation by Polθi varied considerably (SF_ART558_/SF_DMSO_ range: 0.53 to 1.16), with 72% of the cancer cell lines showing substantial radiosensitisation (SF_ART558_/SF_DMSO_<0.9). The average radiosensitisation by Polθi did not differ among the three tumour types (Figure 1B). Moreover, the radiosensitising effect of Polθi did not correlate with *POLQ* expression or the frequency of Polθ-generated DNA scars^7^ –putative indicators of Polθ activity (Figure 1C-D and Supplementary Figure 1A-B). These results highlight the broad potential of Polθi as a radiosensitisation strategy. Nonetheless, the pronounced variability in radiosensitisation, along with the lack or limited response observed in 28% of the models, underscores the need to identify determinants of Polθi-induced radiosensitisation to maximise the clinical benefit of combining Polθi with RT.

**Figure 1.**
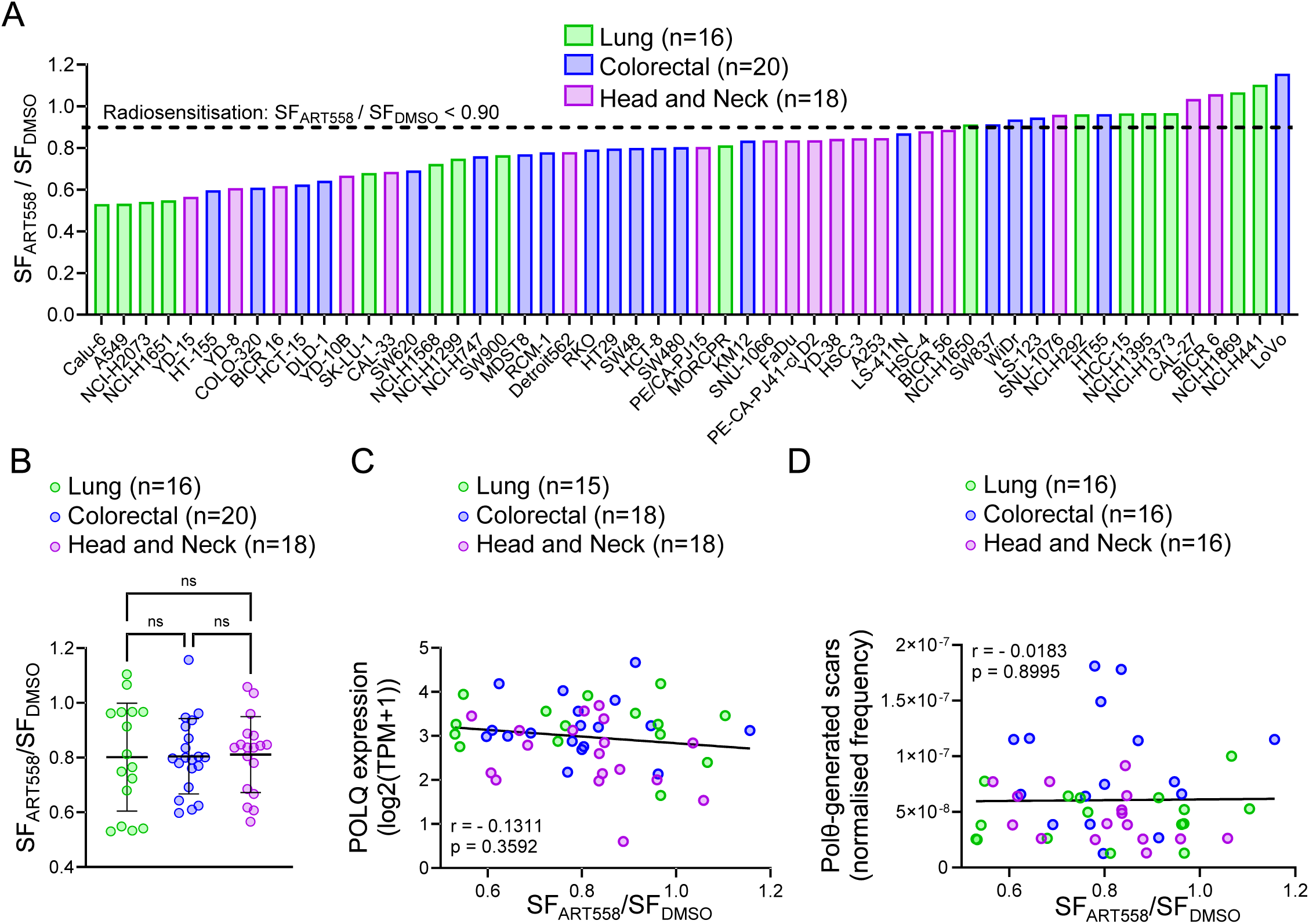
Polθ-induced radiosensitisation varies across cancer cell lines and does not correlate with indicators of Polθ activity (A) Radiosensitisation by **Polθ** in a panel of lung, colorectal and head and neck cancer cell lines, estimated as the surviving fraction (SF) ratios of ART558 to DMSO treated cells (SF_ART558_/SF_DMSO_). (B) SF_ART558_/SF_DMSO_ values stratified by tumour type (mean ± SD). (C) Correlation between *POLQ* expression and the magnitude of radiosensitisation by Polθ (D) Correlation between the frequency of Polθ generated scars per exomic base pair and the magnitude of radiosensitisation by Polθ. Dots represent individual cell lines. Statistical significance in (B) was calculated using one way ANOVA. Correlation coefficients (r) in (C D) were calculated using the Spearman’s method. The black lines in (C D) are regression lines.

### A CRISPR knockout (KO) screen identifies genetic modulators of Polθi-induced radiosensitisation

To identify genetic determinants of the response to Polθi and RT, we conducted a CRISPR KO screen in Cas9-expressing DLD-1 cells targeting 2776 genes involved in the DNA damage response and cancer biology (Figure 2A, Supplementary Figure 2A, Supplementary Data 1A). Screen quality controls confirmed optimal transduction and Cas9 cutting efficiency, adequate representation, and high specificity and sensitivity in the detection of essential genes (Supplementary Figure 2B-E). Using MAGeCK analysis of relative sgRNA abundance at the different timepoints post-treatment, we identified several specific gene KOs conferring significant sensitisation to RT, Polθi, and the combination treatment (Figure 2B-D, Supplementary Figure 2F, and Supplementary Data 1D-E).

**Figure 2.**
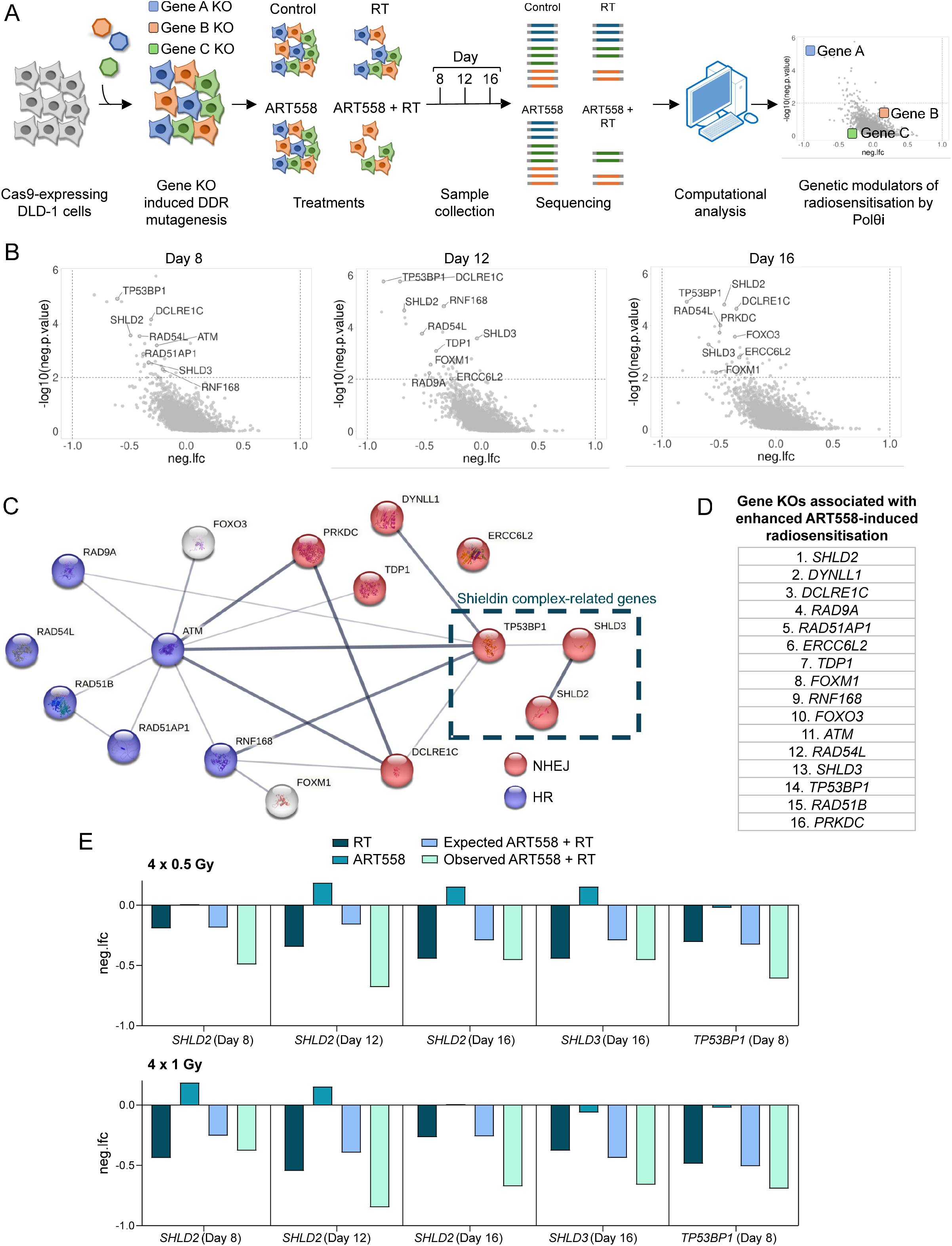
**A CRISPR KO screen identifies genetic modulators of the radiosensitising effect of Polθ**i. (A) Outline of the CRISPR KO screen to identify genetic modulators of Polθi-induced radiosensitisation. The timeline is described in Supplementary Figure 2A. (B) Volcano plots highlighting the specific genes whose knockout synergised with combined Polθi (ART558) and RT (4 x 0.5 Gy) in at least two of the time points and/or RT schedules. For the volcano plots corresponding to the 4 x 1 Gy schedule see Supplementary Figure 2F. neg.lfc: negative log fold change. (C) Network of protein-protein interactions among the genes whose knockout was associated with enhanced Polθi-induced radiosensitisation, generated with the STRING database. The thickness of the connecting lines indicates the strength of the data supporting the interaction. Genes involved in HR and NHEJ have been highlighted in blue and red, respectively. (D) Rank of suppressors of Polθi-induced radiosensitisation identified in the CRISPR screen. (E) Bar graphs representing the neg.lfc values relative to the untreated group for the Shieldin complex-related genes *SHLD2*, *SHLD3,* and *TP53BP1*. Only the timepoints and RT schedules where a meaningful deviation between the RT + ART558 (observed) and the sum of the RT alone and ART558 alone (expected) neg.lfc values (Δ observedf-Expected ≥-0.1) for these genes are shown.

Among the specific gene KOs conferring enhanced sensitisation to the combination of Polθi and RT –exceeding the expected additive effects of either agent alone (see Methods)– we identified factors primarily involved in DSB repair via non-homologous end-joining (NHEJ) and HR (Figure 2B-E, Supplementary Figure 2F and Supplementary Data 1E). This is consistent with the previously reported role of Polθ in driving DSB repair when these pathways are compromised^8,9,13,14^. Remarkably, three physically interacting proteins of the TP53BP1/Shieldin pathway were identified as suppressors of the radiosensitising effect of Polθi: *SHLD2* (*FAM35A*), *SHLD3* (*RINN1*) and *TP53BP1* (Figure 2B-E, Supplementary Figure 2F and Supplementary Data 1E). Of these, *SHLD2* was the top-ranking gene, with its loss consistently showing synergistic negative enrichment in the RT and Polθi treatment arm at the two radiation doses across all experimental timepoints (Figure 2D-E and Supplementary Data 1E).

### *SHLD2* is frequently lost in prostate cancer patients

We next interrogated databases of large prostate cancer cohorts^15–20^ and found that *SHLD2* is frequently lost in prostate cancer, with up to 10% of patients displaying homozygous deletions in this gene (Figure 3A). Notably, approximately a 4% of patients in the TCGA cohort^19,20^—which comprises primary, localised prostate tumours which are typically treated with curative-intent radiotherapy— also exhibited homozygous *SHLD2* deletions (Figure 3A).

**Figure 3.**
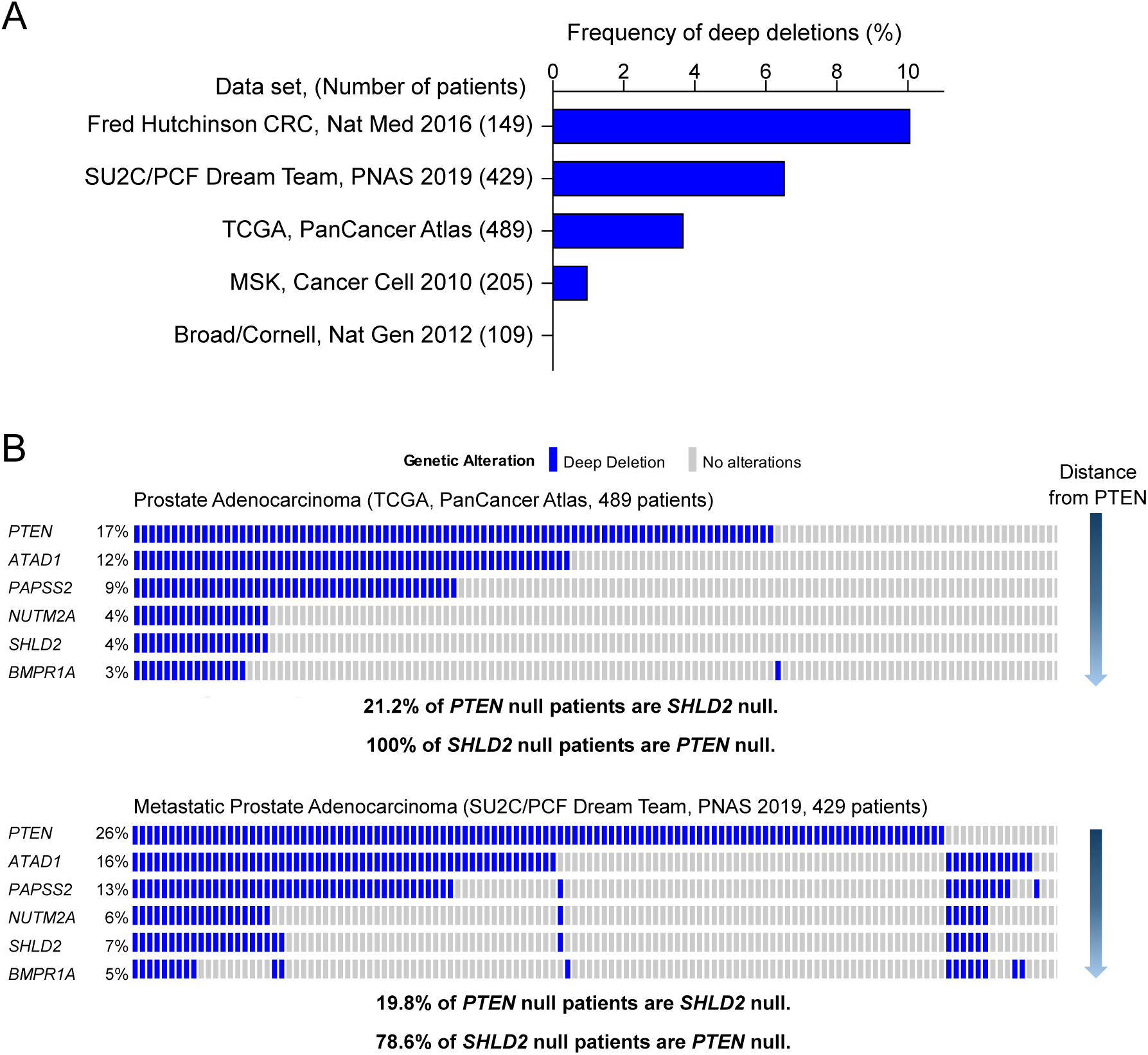
*SHLD2* is frequently lost in human prostate tumours (A) Proportion of tumours with homozygous *SHLD2* deletions in the indicated prostate cancer datasets. Number in brackets indicate the total number of samples with CNV data available. (B) Proportions of homozygous deletions in *PTEN* and neighbour genes. Each bar within the same column represents the same patient, enabling visualisation of co-occurring deletions.

*SHLD2* is located less than 1 Mbp away from *PTEN* (Supplementary Figure 3), a tumour suppressor gene that is frequently deleted in prostate cancer and whose loss is associated with adverse outcomes^21^. We found that homozygous *SHLD2* deletions occur in approximately ∼20% of tumours with homozygous *PTEN* deletions, and that between 79-100% of tumours with homozygous *SHLD2* deletions harbour homozygous *PTEN* deletions (Figure 3B). In addition, we observed that co-deletion rates among genes flanking *PTEN* decrease with increasing distance from *PTEN* (Figure 3B and Supplementary Figure 3). Taken together, these findings indicate that *SHLD2* loss may constitute a collateral therapeutic vulnerability in prostate cancer that can be exploited through the combination of Polθi and RT.

### *SHLD2* loss synergises with combined Polθi and radiotherapy *in vitro*

Loss of TP53BP1/Shieldin has previously been shown to confer PARPi resistance and to synergise with Polθi monotherapy specifically in HR-deficient cells^10,12^. It is important to note that the DLD-1 cells used in our CRISPR KO screen are HR-proficient^10^. Therefore, our observation that TP53BP1/Shieldin deficiency may sensitise HR-proficient cells to combined Polθi and RT represents a novel finding.

Given the clinical relevance of *SHLD2* loss in prostate cancer, we generated *SHLD2* KO clones (*SHLD2*^-/-^) from the HR-proficient^22^ and *PTEN*-expressing^23^ prostate cancer cell lines DU145 (clone A2) and 22Rv1 (clones A6, C1, and G6) (Supplementary Figure 4A), to validate *SHLD2* loss as both a therapeutic vulnerability and a potential predictive biomarker of Polθi-mediated radiosensitisation. Since, according to our co-deletion analysis (Figure 3B), most prostate cancer tumours with homozygous *SHLD2* deletions are also *PTEN*-null, we knocked out *SHLD2* in the *PTEN*-null^24^, HR-proficient^22^ breast cancer cell line CAL-51 (clones E1 and E2) (Supplementary Figure 4A), to rule out an effect of *PTEN* loss on Polθi-induced radiosensitisation. Compared to their parental counterparts, the *SHLD2*^-/-^ clones were more sensitive to RT alone *in vitro* (Figure 4A-C), in line with the role of the TP53BP1/Shieldin pathway in promoting DSB repair by NHEJ^25–27^ and with the results of our screen (Supplementary Data 1D). Continuous treatment with ART558 alone for 14 days did not affect the colony-forming ability of the *SHLD2*^-/-^ cells (Figure 4D-F), confirming that these HR-proficient models are not susceptible to Polθi monotherapy, unlike what was reported previously for HR-deficient, TP53BP1/Shieldin-deficient cells^10,12^. However, compared to the *SHLD2*^WT^ parental lines, the extent of Polθi-induced radiosensitisation with ART558 was significantly higher in the *SHLD2*^-/-^ clones, irrespective of the *PTEN* status (Figure 4G-J). Reconstitution of *SHLD2* expression in the 22Rv1 *SHLD2*^-/-^ clones reversed the radiosensitising effect of Polθi, confirming the direct role of *SHLD2* in modulating this response (Figure 4K-L and Supplementary Figure 4B).

**Figure 4.**
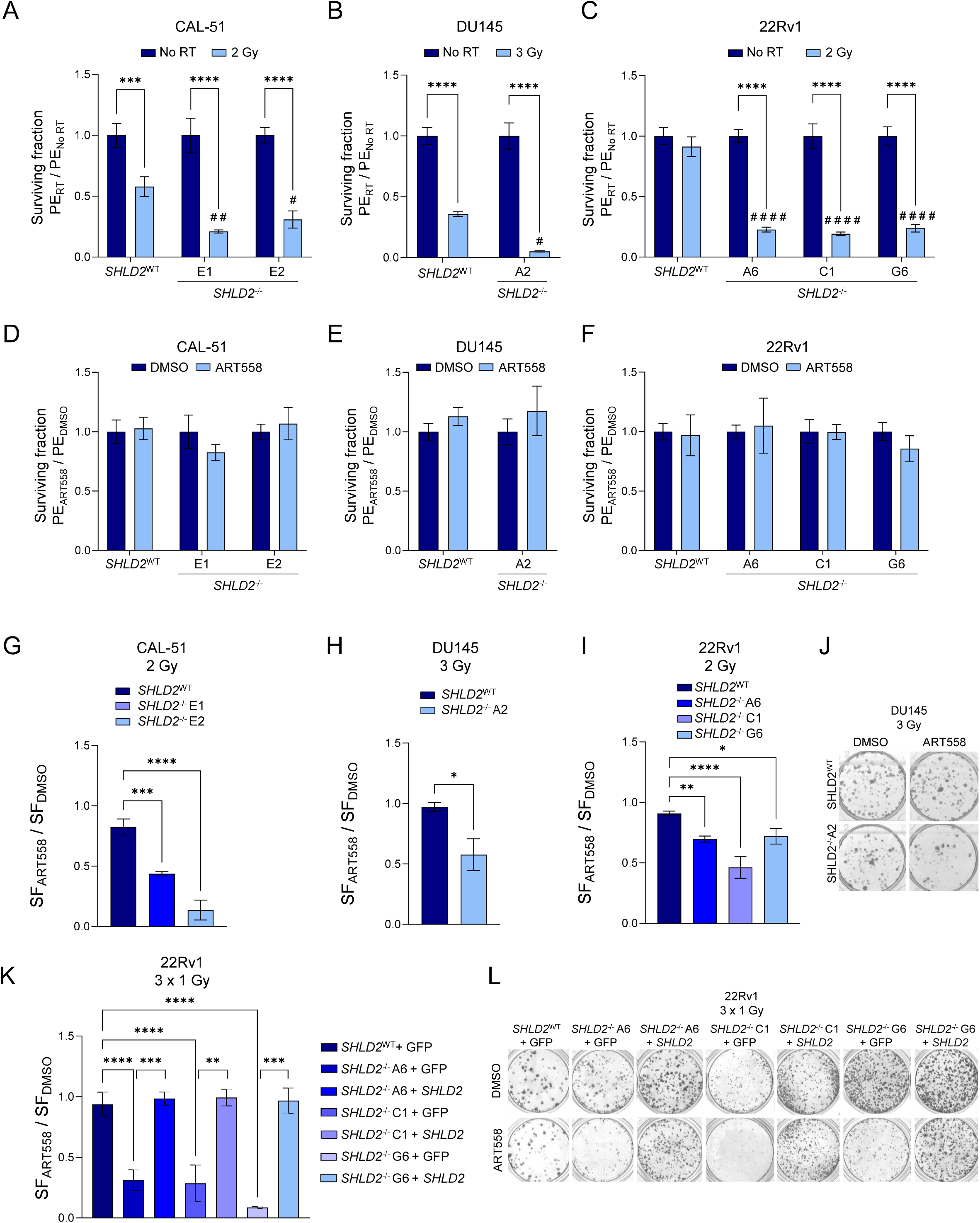
***SHLD2*** loss synergises with combined Pol8i and RT ***in vitro*** Colony-forming capacity of *SHLD2*^WT^ *vs SHLD2*^-/-^ CAL-51, DU145 and 22Rv1 cells treated with: (A-C) RT alone, normalised to unirradiated controls. (D-F) 4 μegd/L Polθi ART558 alone for 14 days, normalised to DMSO-treated control. (G-F) 4 μmol/L polθi ART558 and RT, expressed as the surviving fraction (SF) ratios of ART558-to DMSO-treated cells, to show the radiosensitising effect of Polθi. (J) Representative colony formation assays of *SHLD2*^WT^ and *SHLD2*^-/-^ DU145 cells treated with RT and either DMSO or ART558. (K) Colony formation capacity of the *SHLD2*^WT^ *vs SHLD2*^-/-^ 22Rv1 models transduced with either a vector expressing *SHLD2* or a GFP control vector. Radiosensitisation by Polθi is shown as the surviving fraction (SF) ratios of ART558-to DMSO-treated cells. (L) Representative colony formation assays from (K). Mean ±SD of three technical replicates (representative of two independent experiments). Statistical tests: In (A-C) Two-way ANOVA and Tukey’s post-hoc test; hashes and asterisks refer to the comparison of RT-treated *SHLD2*^WT^ *vs* RT-treated *SHLD2*^-/-^ clones, and the comparison between the No RT and RT-treated samples for each indicated clone, respectively. # or *p<0.05; # # or **p<0.01; # # # or ***p< 0.001; # # # # or ****p< 0.0001. (D-F) Multiple unpaired t-tests; (G-I, K) One-way ANOVA and Dunnett’s post-hoc test for CAL-51 and 22Rv1 graphs; Welch’s t-test for DU145 graphs. *p<0.05; **p<0.01; ***p< 0.001; ****p< 0.0001.

### *SHLD2*-deficient cells are more dependent on Polθ to prevent DSB accumulation and genomic instability following RT

To investigate the mechanism by which *SHLD2* loss confers sensitivity to Polθi and RT, we assessed the dynamics of DSB markers following RT in DU145 *SHLD2*^WT^ and *SHLD2*^-/-^ A2 cells, using fluorescence microscopy. Compared to the *SHLD2*^WT^ cells, the *SHLD2*^-/-^ cells exhibited increased levels of γH2AX, 53BP1, pATM and RAD51 nuclear foci, becoming significant as early as 3-6 hours post-RT (Figure 5A-J). Importantly, a further increase in these DSB markers was consistently observed upon Polθi treatment at 24-48 hours after RT in the *SHLD2*^-/^*^-^* A2 DU145 cells, but not in their parental *SHLD2*^WT^ counterparts (Figure 5A-J). We also determined the induction of cytoplasmic micronuclei in the *SHLD2*^WT^ and *SHLD2*^-/-^ A2 DU145 cell lines after RT, as a marker of genomic instability^28^. In line with the DSB foci data, the proportion of RT-induced micronuclei was higher in the *SHLD2*^-/-^ cells than in the parental cells (Figure 5K-L). Moreover, further induction of micronuclei after RT was observed only in the *SHLD2*^-/-^ cells treated with Polθi (Figure 5K-L). These results demonstrate that *SHLD2*-deficient cells are more reliant on Polθ to prevent RT-induced DSB accumulation and chromosomal instability.

**Figure 5.**
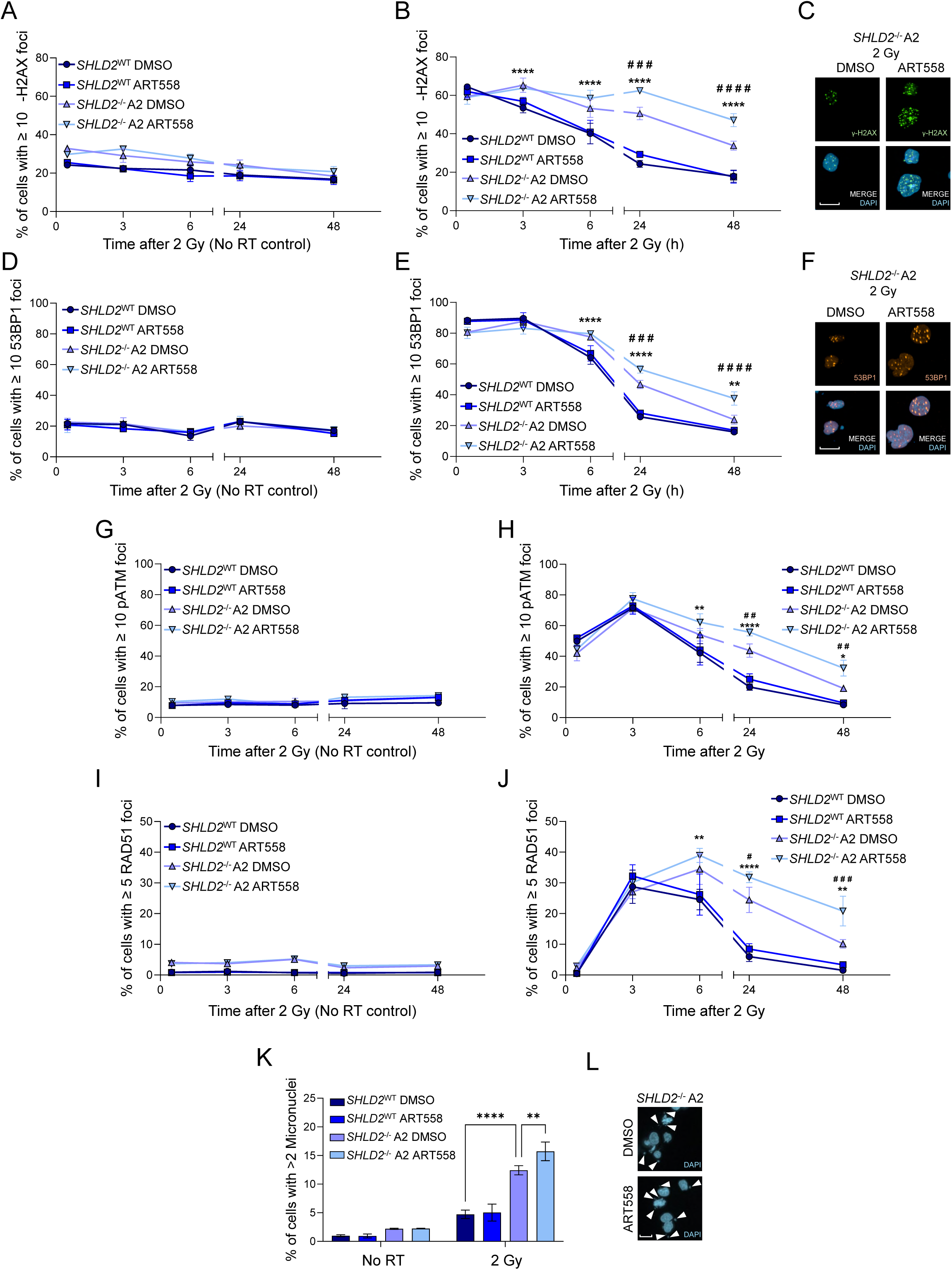
*SHLD2*^-/-^ cells rely more on Pol8 to prevent RT-induced DSB accumulation and chromosomal instability. *SHLD2*^WT^ and *SHLD2*^-/-^ A2 DU145 cells were treated with 4 μmol/L Polθi ART558 and the indicated RT doses. (A, B) yH2AX foci dynamics for unirradiated and irradiated conditions respectively, assessed by fluorescence microscopy. (C) Images from (B) at 24 h post-RT. (D, E) 53BP1 foci dynamics for unirradiated and irradiated conditions respectively (F) Images from (E) at 24 h post-RT. (G, H) pATM foci dynamics for unirradiated and irradiated conditions respectively (I, J) RAD51 foci dynamics for unirradiated and irradiated conditions respectively (K) Micronuclei formation 48 h after RT, assessed by fluorescence microscopy. (L) Images of RT-treated *SHLD2*^-/-^ A2 cells from (K). Arrowheads indicate micronuclei. Data are representative of three independent experiments, showing mean ± SD from triplicate wells. Statistical significance was calculated with two-way ANOVA and Tukey’s post-hoc test. In (B, E, H, J) hashes and asterisks refer to the comparison of DMSO-treated *SHLD2*^-/-^ *vs* ART558-treated *SHLD2*^-/-^ cells, and DMSO-treated *SHLD2*^WT^ *vs* DMSO-treated *SHLD2*^-/-^ cells, respectively. # or *p<0.05; # # or **p<0.01; # # # or ***p< 0.001; # # # # or ****p< 0.0001. Scale bars in (C, F, L) = 20µm.

*SHLD2*-deficient tumours are highly sensitive to combined Polθi and radiation treatment *in vivo*.

We next assessed the effect of Polθi combined with fractionated RT in NRG mice inoculated with the DU145 *SHLD2*^WT^ and *SHLD2*^-/-^ A2 cells (Figure 6A). In this experiment, we used ART899, an analogue of ART558 previously shown to display improved pharmaceutical properties, making it suitable for *in vivo* studies^2^. Importantly, treatment with the Polθi ART899 in combination with fractionated RT was well tolerated, as indicated by the absence of significant changes in body weight (Figure 6B). Consistent with our *in vitro* data, ART899 monotherapy did not affect the growth of either *SHLD2*^WT^ or *SHLD2*^-/-^ tumours, and the *SHLD2*^-/-^ tumours were more sensitive to RT alone than their *SHLD2*^WT^ counterparts (Figure 6C-E). Importantly, compared to RT alone, the addition of ART899 to RT induced a significant reduction in tumour growth and significant extension of survival in the *SHLD2*^-/-^ xenografts, but not in *SHLD2*^WT^ xenografts (Figure 6C-E). Overall, our results demonstrate that *SHLD2* loss renders tumours more susceptible to the combination of Polθi with RT *in vivo*.

**Figure 6.**
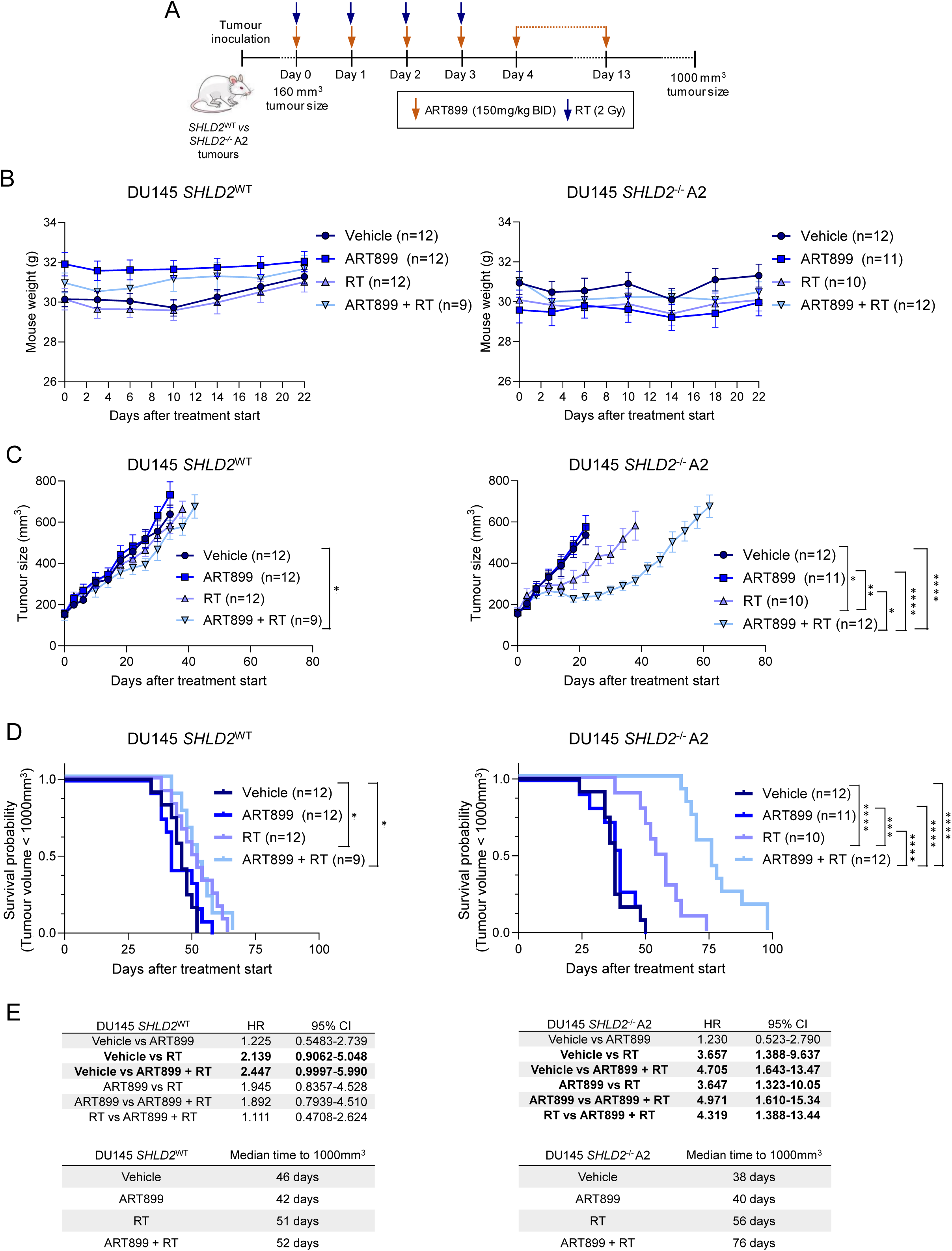
*SHLD2*-deficient tumours are highly sensitive to combined Pol8i and radiation treatment *in vivo*. *SHLD2^WT^* and *SHLD2^-/-^* A2 DU145 xenografts in NRG mice treated with 4 x 2 Gy X-Ray (RT) and/or the Polθi ART899 (150 mg/kg twice daily for 14 days). (A) Diagram of the experimental design. (B) Mouse weights. (C) Tumour growth kinetics of *SHLD2^WT^ and SHLD2^-/-^ A2* DU145 xenografts. Mean ± SE. Unpaired Mann-Whitney U-tests were used to assess differences in tumour size averages at Day 22. *p<0.05; **p<0.005; *** p<0.0005 ; ****p<0.0001 (D) Kaplan-Meier survival plot for a tumour threshold of 1000mm^3^. Log-rank (Mantel-Cox) test: *p<0.05; **p<0.005; *** p<0.0005 ; ****p<0.0001. (E) Tables show hazard ratios (HR) and 95% confidence intervals (CI) for the indicated comparisons (Mantel-Cox test), and median survival times for each treatment group.

## DISCUSSION

In the present study, using a comprehensive panel of cancer cell lines, we demonstrate that Polθi exerts a radiosensitising effect in approximately three quarters of the models analysed, underscoring the broad therapeutic potential of combining Polθi with RT. However, the extent of Polθi-induced radiosensitisation varied considerably across the 54 cancer cell lines screened, with a non-negligible subset of models exhibiting minimal or no response. This variability underscores the need for biomarkers to identify patients most likely to benefit from Polθi in combination with RT, which would significantly ease clinical translation and, ultimately, improve patient outcomes. The lack of correlation between indicators of Polθ activity and the extent of Polθi-induced radiosensitisation prompted us to conduct a forward genetic CRISPR KO screen to uncover determinants of Polθi-induced radiosensitisation.

In our CRISPR KO screen, components of the TP53BP1/Shieldin pathway (*SHLD2*, *SHLD3* and *TP53BP1*) were among the genes whose loss was associated with enhanced radiosensitisation by Polθi, with *SHLD2* as the highest-ranked hit. Here, we demonstrate that *SHLD2* loss markedly enhances tumour cell sensitivity to combined Polθi and RT in both cancer cell lines *in vitro* and in tumour xenografts. This finding is clinically relevant, as we identified a subset of prostate cancer patients with homozygous *SHLD2* deletions, defining a patient population that could extensively benefit from Polθi combined with RT, and offering a potential biomarker for patient stratification.

We found that ∼20% of *PTEN*-null prostate tumours also harbour homozygous deletions in *SHLD2*, and that the majority of *SHLD2*-null tumours are likewise *PTEN*-null. The inverse correlation between the distance of neighbouring genes from *PTEN* and their co-deletion incidence suggests that *PTEN* is the tumour suppressor acting primarily as a cancer driver, while the deletion in the nearby genes –including *SHLD2*– are likely passenger deletion events. *PTEN* loss has been correlated with adverse outcomes in prostate cancer patients, including those treated with RT^21,29–32^. We demonstrate here that the enhanced radiosensitising effect of Polθi is preserved in *SHLD2*-deficient cells with *PTEN* loss, indicating that *PTEN* deficiency does not compromise the response to Polθi. Therefore, *SHLD2* loss may be a collateral vulnerability to Polθi following RT in *PTEN*-deficient tumours, offering a strategy to effectively target a subset of this group of patients.

Of note, we also demonstrated that *SHLD2* loss sensitises cancer cells to RT alone both *in vitro* and *in vivo*. This finding raises the possibility that patients with *PTEN*-deficient/*SHLD2*-deficient tumours may experience better outcomes after RT than those with *PTEN*-deficient/*SHLD2*-proficient tumours. However, we were unable to test this hypothesis due to the limited number of RT-treated patients with available clinical outcome data in the publicly available cohorts analysed. Future studies should address how *SHLD2* loss influences prognosis both in *PTEN*-proficient and -deficient patient populations, to better gauge the potential clinical impact of combining Polθi with RT in these subgroups.

Genetic or pharmacological inhibition of Polθ previously revealed a synthetically lethal interaction between Polθ and components of the HR repair pathway, such as *FANCD2*, *BRCA1* or *BRCA2*^8–10^. Depletion of components of the TP53BP1/Shieldin pathway has also been shown to sensitise cancer cells to Polθi monotherapy specifically in HR-deficient backgrounds^10^. In contrast, the *SHLD2*^-/-^ models used in our study are HR proficient, underscoring the potential utility of Polθi beyond HR-deficient tumours. In line with our earlier reports^2^, the present study confirms that the addition of Polθi to fractionated RT is well tolerated *in vivo.* Notably, several Polθ inhibitors –including the class of compounds tested in this study– have now entered early-phase clinical trials (NCT05898399, NCT04991480, NCT06077877, NCT05687110, NCT06666270, NCT06545942, NCT06560632), further supporting the translational promise of this combination strategy.

Among the other hits whose loss conferred enhanced sensitisation to the combination of Polθi and RT, we identified several genes involved in the major DSB repair pathways, namely HR and NHEJ. This is in agreement with the role of Polθ-mediated repair of DSBs as a backup mechanism to HR and NHEJ^8,9,13,14^. Although loss or inactivation of some of these genes like *ERCC6L2*^33^, *PRKDC* (*DNA-PK*)^34,35^ and *ATM*^36^ has been shown to sensitise cells to RT alone, this is the first report identifying them as potential modulators of Polθi-induced radiosensitisation. Further studies are warranted to evaluate their relevance for patient selection. It is tempting to speculate that available inhibitors targeting some of these proteins –such as ATM or DNA-PK– could potentiate the radiosensitising effect of Polθi. However, clinical translation of these strategies would have to face the complexity of triple combination therapies and the toxicity of these inhibitors when combined with RT^37,38^. Our mechanistic experiments suggest that, upon *SHLD2* loss, cells become more dependent on Polθ to prevent DSB accumulation and maintain genome integrity following RT exposure, which explains the enhanced radiosensitisation observed with Polθi in *SHLD2*-deficient cells.

In summary, we demonstrate that *SHLD2* loss increases the susceptibility of cancer cells to the radiosensitising effect of Polθi. *SHLD2* loss may serve as a biomarker for selecting prostate cancer patients most likely to benefit from Polθi in combination with RT. This vulnerability reflects the increased dependence of *SHLD2*-deficient cells on Polθ to prevent DSB accumulation and maintain chromosomal stability. These findings support the clinical investigation of this combination strategy for patients with *SHLD2*-deficient tumours.

## METHODS

### Cell culture and treatments

Cell lines were cultured in a humidified incubator at 37 °C and 5% CO_2_, passaged before reaching 80% confluency and were routinely tested for mycoplasma using the MycoAlert PLUS Mycoplasma Detection kit (Lonza). Details of cell lines, their source and cell culture media are provided in Supplementary Table 1. ART558 was produced as described previously^10^. ART899 was produced from its non-deuterated form, ART812, with the addition of deuterium during synthesis, as previously described^10^. Compounds were stored in powder form in a vacuum chamber at room temperature in the dark. Compounds were reconstituted in DMSO at 12 mmol/L and these stocks were kept at room temperature in the dark. ART558 was added one hour prior to RT.

### Cell line panel experiment

We used a panel comprising 16 lung, 20 colorectal, and 18 head and neck cancer cell lines (listed in Figure 1 and Supplementary Table 1). Cells were seeded in 24 well-plates at optimised seeding densities. The day after seeding, cells were treated with 3 µmol/L ART558 and exposed to the first radiation fraction of a 3 x 2 Gy regimen; subsequent fractions where delivered every 24 h. The inhibitor was maintained throughout the experiment. Cells were incubated for 14 days post-treatment and then stained with crystal violet. Cell viability in the cell line panel was quantified using crystal violet solubilisation, since many of the cell lines used are not capable to grow detectable colonies. Stained samples were incubated for at least 30 minutes with 10% acetic acid to solubilise the crystal violet dye. After complete solubilisation, the optical density at 590 nm of the acetic acid solution was measured using a Clariostar plate reader. Absorbance values were then normalised to the seeding numbers, and surviving fractions (SF) were calculated as the corrected absorbance of the irradiated condition divided by that of the unirradiated control. The extent of radiosensitisation by Polθi was estimated as SF_ART558_/ SF_DMSO_. The SF_ART558_/ SF_DMSO_ values were then correlated with POLQ expression levels and the frequency of exomic Polθ-generated scars. POLQ expression values were retrieved from the DepMap Public 24Q4 RNA seq dataset^39^. To quantify the frequency of exomic Polθ-generated scars –defined here as deletions of ≥5 bp flanked by microhomologies of ≥2 bp– we applied the computational pipeline described by Carvajal-Garcia *et al.*^7^ to the DepMap 24Q4 whole exome sequencing (WES) dataset^39^. Scar frequencies were normalised for each cell line to their total exonic length in base-pairs, calculated as the sum of Ensembl-annotated exon lengths weighted by copy number at the gene level retrieved from DepMap.

### CRISPR KO Screen

We generated a Cas9-expressing cell line (DLD-1 Cas9) by infecting DLD-1 cells with LVCAS9BST-1EA lentiviral particles carrying a vector encoding for blasticidin resistance and Cas9 genes^40^. After infection, cells were treated with 10 μg/mL blasticidin for 10 days. Optimal (>95%) Cas9 cutting efficiency in Cas9-expressing DLD-1 cells was demonstrated using the CRISPRuTest™ (Cellecta) (Supplementary Figure 2B-C).

For the CRISPR KO screen, we utilised a pooled lentiviral CRISPR KO library targeting genes implicated in the DNA damage response and in cancer pathogenesis (Supplementary Data 1A). This library comprised 16732 gRNAs targeting 2776 genes, with approximately six sgRNAs per gene, along with 100 non-targeting control sgRNAs. Fifty million DLD-1 Cas9 cells were infected with this library at a multiplicity of infection (MOI) of 0.4, and incubated for 7 days with 2 μg/mL puromycin to select for cells with successful donor sequence integration.

The experimental design and assay timeline are outlined in Supplementary Figure 2A. Library-infected cells were seeded one day before treatment, with triplicates per treatment group. A minimum of 13 million library-infected cells were used for each replicate and cultured throughout the screen to maintain a library representation of at least 760-fold. Treatment groups were as follows: DMSO (vehicle control), 6 μmol/L ART558, 0.5 Gy, 1 Gy, 0.5 Gy + 6 μmol/L ART558 and 1 Gy + 6 μmol/L ART558. For RT treatment, cells received four fractions of 0.5 or 1 Gy every 24 h using a CellRad (Faxitron®) X-ray irradiator at a dose rate of 1.02 Gy/min. ART558 was added 1 hour before the first RT dose.

On Day 4, cells were detached 3 hours post-RT, counted, and re-seeded to maintain exponential cell growth and library representation. At this point, 6 μmol/L ART558 was added again and kept in the media until collection of cells. Cell pellets were collected for DNA extraction on days 8, 11 and 15 after treatment initiation. The growth curves of cells exposed to the different treatments over the course of the experiment are displayed in Supplementary Figure 2D.

Genomic DNA (gDNA) extraction and purification was carried out using GenElute™ Mammalian Genomic DNA Miniprep Kit (G1N70-1KT, Sigma-Aldrich), and gDNA solutions were prepared at a 100 ng/µL concentration for PCR. sgRNA sequences integrated in the gDNA were recovered by nested PCR amplification utilising vector-backbone-directed primers. Parallel PCRs were performed with a Veriti™ 96-well thermal cycler (Applied Biosystems™) under the following conditions: PCR1: 95°C for 10s, 38 cycles of 95°C for 10s, 55°C for 10s, 72°C for 20s, 72°C for 1 min; PCR2: 95°c for 10s, 12 cycles of 95°C for 10s, 56°C for 10s, 72°C for 20s, 72°C for 1 min; PCR3: 95°c for 10s, 12 cycles of 95°C for 10s, 50°C for 10s, 72°C for 22s, 72°C for 1 min.

PCR products from parallel reactions were subsequently pooled and purified using GenElute™ PCR Clean-Up Kit (NA1020-1KT, Sigma-Aldrich). Sequencing reactions were performed on an Illumina NextSeq 500/550 using the High Output v2 kit, following the manufacturer’s instructions. The sequencing reads were computationally demultiplexed and trimmed to remove barcodes and adapters. The trimmed reads were aligned to the reference genome with BowTie2^41^, filtering out ambiguous hits. SAMtools^42^, was then applied to process the alignment files and generate depth data for the library sequences. Z-scores for each sequence was calculated to provide a normalised dataset.

The sgRNA counts obtained as described above were analysed with MAGeCK^43^ to generate gene-level enrichment scores for each of the treatment groups in relation to the corresponding control group (Supplementary Data 1C). Screen quality was assessed at both sgRNA and gene levels. The Receiver Operating Characteristic (ROC) curve was generated using “AchillesCommonEssentialControls.csv” and “AchillesNonessentialControls.csv” from DepMap version 22Q4, and used to define essential and non-essential genes, respectively. This analysis showed high specificity and sensitivity of essential genes, utilised here as positive controls (Supplementary Figure 2E).

A negative sgRNA enrichment for a given gene in the ART558, RT, or ART558 + RT groups was considered significant when that gene exhibited a MAGeCK negative selection score (neg.score) ≤ 0.001 and a negative selection p-value (neg.p.value) < 0.01 in at least two conditions within those groups –i.e., across the three collection time points and/or, for RT-treated samples, across the two radiation regimens. Genes meeting these thresholds were ranked by the sum of the log fold changes (∑neg.logFC) of all qualifying conditions (i.e., those time points or radiation regimens in which neg.score ≤ 0.001 and neg.p-value < 0.01). This yielded a list of the candidate genes whose knockout was associated with enhanced sensitivity to either ART558 or RT (Supplementary Data 1D).

We then generated a list of genes whose KO was associated with enhanced radiosensitisation by Polθi, with an effect above the expected additivity of the two single treatments, calculated using the Bliss independence model^44^. To this aim, we first selected the hits fulfilling the above mentioned criteria of neg.score ≤ 0.001 and neg.p.value < 0.01 for ART558 + RT. Then, we determined the expected combined neg.lfc (E_RT+ART558_) by summing the individual observed (O) effects of RT and ART558 alone according to the following equation: E_RT+ART558_ = O_RT_(neg.lfc) + O_ART558_(neg.lfc). Then, the deviation between the observed neg.lfc values for the combination treatment (O_RT+ART558_) and E_RT+ART558_ was calculated as: ΔO−E= O_RT+ART558_− E_RT+ART558_. Finally, a specific gene KO was classified as a genetic vulnerability to the ART558 + RT combination when ΔO−E≤−0.1 in at least two of the six ART558 + RT treatment conditions. For each gene, the neg.lfc scores across the qualifying treatment conditions (ΔO−E≤−0.1) were summed to yield an overall ∑(ΔO−E) score. Genes were then ranked by descending ∑(ΔO−E) (Figure 2D and Supplementary Data 1E).

### Protein interaction network analysis

STRING portal (https://string-db.org/) was used to generate the protein-protein interaction network of the candidate modulators of Polθi-mediated radiosensitisation identified in the CRISPR KO screen.

Interrogation of cBioportal for frequency of *SHLD2* loss in clinical samples.

A non-redundant set of prostate studies with gene copy number data for over 100 patients was selected on cBioPortal and queried for *SHLD2* homozygous deletions (SHLD2: HOMDEL). For the two representative studies TCGA (PanCancer Atlas)^19,20^ and SU2C/PCF^17^ Dream Team that provided data amenable for co-deletion representation, homozygous deletions were plotted for the main genes in the genomic region between *PTEN* and *SHLD2* to assess the frequency and co-occurrence pattern of homozygous deletions (deep deletions defined as per <u>cBioportal)</u>.

The results pertaining to the TCGA (PanCancer Atlas)^19,20^ are based upon data generated by the TCGA Research Network: https://www.cancer.gov/tcga.

### Colony formation assays (CFAs)

Cells were seeded in either 24-well or 6-well plates and allowed to settle for 4-5 hours. 4 μmol/L of ART558 was added to each well with DMSO utilised as a negative control. An X-ray irradiator (CellRad by Faxitron®; dose rate 1.02 Gy/min) was used to irradiate CFA. Media containing ART558 or DMSO was left for the duration of the assay.

The plates were allowed to form colonies and then stained and fixed with crystal violet 12-14 days later. Colonies were analysed using an Oxford Optronics Gel Count machine and GelCount software. The plating efficiency (PE) was defined as: PE = average colony number/cells plated. In the experiments testing the single-agent effect of ART558 (Figure 4D-F), the surviving fraction (SF) relative to the DMSO control was calculated as PE_ART558_/PE_DMSO._ In the experiments involving RT treatment, the SF relative to the unirradiated control (No RT) was calculated as PE_RT_/PE_No RT_. To estimate the extent of radiosensitisation by ART558, we calculated the ratio of the SFs of ART558-treated to DMSO-treated cells.

### Immunofluorescence microscopy

Cells were seeded in black-wall 96 well-plates and left to attach overnight. The following day, cells were treated with 4 μmol/L ART558. An X-ray irradiator (CellRad by Faxitron®; dose rate 1.02 Gy/min) was used to irradiate the plates. At the corresponding timepoints after treatment, cells were washed with PBS and pre-extracted with CSK buffer (Supplementary Table 2) for 6 minutes. Cells were then fixed with 4% PFA in PBS for 10 minutes. Samples were incubated for 1 hour at room temperature with blocking buffer (0.5% BSA, 0.5% Triton in PBS), then with primary antibodies (see Supplementary Table 3) diluted in blocking buffer overnight at 4°C, and then with the secondary fluorescent antibodies (see Supplementary Table 3) for one hour at room temperature in the dark. Samples were imaged using an Operetta CLS system, and processed and analysed on the Harmony image analysis software. An analysis pipeline was generated and applied to all acquired images in the Harmony software to quantify foci and micronuclei.

### Generation of *SHLD2*-deficient clones, knockout confirmation, and *SHLD2* restoration

The *SHLD2*^-/-^ CAL-51 clones E1 and E2, DU145 clone A2, and 22Rv1 clones A6, C1 and G6, were generated by Oxford Genetics (Oxford, UK), using CRISPR/Cas9 editing as described previously^10^. Briefly, synthetic guide RNAs (sgRNA) for CRISPR/Cas9 were designed to target the *SHLD2* gene (reference gene ENSG00000122376). The 5′→3′ sequence of the genomic sgRNA target used was: ATTACAGCATCTCAGAAGAT. Pools of cells carrying the edited gene were generated by transient co-transfection of the sgRNA complexed with CRISPR/Cas9 protein. Single cells were isolated, and the targeted exon was sequenced by Sanger sequencing. Selected clones with out-of-frame insertion/deletions in all alleles were expanded and validated by PCR followed by high-throughput sequencing, which confirmed that all clones used in the study carried out-of-frame insertions or deletions in all *SHLD2* alleles (Supplementary Figure 4A). During the course of the experiments, representative clones were further checked by PCR, Sanger sequencing, and ICE analysis to validate that the *SHLD2* KO status was maintained *in vitro* and *in vivo* (Supplementary Figure 5). More methodological details on the knockout confirmation are provided in Supplementary Methods. To restore *SHLD2* expression in 22Rv1 *SHLD2*^-/-^ clones, cells were infected with a lentiviral vector carrying either the full open reading frame (ORF) of *SHLD2* or the control ORF of GFP (Ex-A8388-Lv105 and Ex-EGFP-Lv105 from Labomics, respectively), as previously described^10^. Infected cells underwent puromycin selection and expansion, and *SHLD2* restoration was confirmed by Western blotting (Supplementary Figure 4B).

### *In vivo* experiments

The project licence covering the animal work was approved by the Animal Welfare and Ethical Review Body (AWERB) at the University of Oxford, and granted by the UK Home Office Animals in Science Regulation Unit (ASRU) under the Animals (Scientific Procedures) Act 1986 (ASPA). We used male NOD.Cg-Rag1^tm1Mom^ Il2^rgtm1Wjl^/SzJ (NRG) mice, aged 7-8 weeks. Mice were subcutaneously injected in the right flank with 2.5 x 10^6^ DU145 *SHLD2*^WT^ or *SHLD2*^-/-^ clone A2 cells suspended in a 1:1 mixture of Matrigel and PBS, in a total volume of 100 µL. Following tumour cell injection, mice were randomly assigned via a random number generator (random.org) to one of four treatment groups: Vehicle control, ART899, RT or ART899 combined with RT. The vehicle consisted of 5% DMSO, 5% ethanol, 20% TPGS and 30% PEG400, diluted in ultrapure water. Mice received either vehicle alone or ART899 (150 mg/kg body weight) dissolved in vehicle, administered by oral gavage twice daily for 14 days (days 0-13), at a volume of 0.043 mL per 10 g body weight. The first daily gavage dose was given 1-2 h prior to RT, and the second dose was administered 6 hours after the first dose. RT was delivered to tumours using a Gulmay 320 X-ray generator (Gulmay Medical Ltd, Camberley, UK) at 300 kV and 10 mA. The fractionated RT regimen consisted of four consecutive doses of 2 Gy delivered every 24 h, on days 0 to 3, under isoflurane anaesthesia. During irradiation, mice were covered with a lead shield that exposed only the tumour to the X-rays.

Treatments started when tumours reached an approximate size of 160 ± 4 mm^3^ (mean ± standard deviation). Animals were weighed before gavage dosing during treatment, and twice weekly thereafter. Tumour size was assessed at least two times a week with callipers and calculated according to the formula: length × width^2^/2. Mice were euthanised with pentobarbital when tumours reached 1000 mm^3^. In the graphs showing tumour size vs days after treatment start, curves were represented up to the time point where the first mouse in the corresponding group reached this tumour size threshold. Survival fractions at specific time points after treatment initiation were estimated based on the number of mice that reached a tumour size of 1000 mm^3^ relative to the initial group size, using the Kaplan-Meier method. Statistical significance of the difference between survival curves was assessed using the log-rank (Mantel-Cox) test.

### Statistical analysis

The specific statistical analyses are indicated in the figure legends and, where relevant, detailed in the corresponding methods section. All graphs were generated and statistical analyses were performed with GraphPad Prism, unless otherwise specified.

## DATA AVAILABILITY

The datasets generated during and/or analysed during the current study are available from the corresponding authors upon reasonable request.

## AUTHOR CONTRIBUTIONS

**G.R.B**: Conceptualisation, supervision, formal analysis, investigation, visualisation, methodology, writing–original draft, writing–review and editing. **P.T:** Conceptualisation, formal analysis, investigation, visualisation, methodology, writing– original draft, writing–review and editing. **Al.Ci:** Contributed to IF experiments, contributed to manuscript figures. **N.M:** Contributed to in vivo experiments. **C.G:** Contributed to in vivo experiments. **D.N:** Contributed to in vitro experiments. **W.C.C:** Bioinformatics data analysis and interpretation. **M.L.B:** Performed and analysed experiments **Au.Ce:** Designed and performed CRISPR screening experiments, data analysis **V.G:** overview of CRISPR screen. **B.R.H:** Contributed to CRISPR screening experiments and data analysis. **K.K:** Bioinformatics data analysis and interpretation. **S.A.L:** Performed and analysed in vitro experiments. **J.B.M:** Designed and supervised in vivo experiments. **Su.Me:** Supervision of bioinformatics data analysis and interpretation. **Sh.Mo:** Planned and performed experiments, data analysis. **J.N:** Performed CRISPR screening experiments, data analysis **N.M.P.D:** Contributed to data generation for the panels of cell lines. **E.R:** Review and interpretation of DDR data. **M.R.L:** Designed and supervised in vivo experiments. **A.T:** Contributed to IF experiments. **R.A.H:** Developed Polθ inhibitors. **G.C.M.S:** Shaping content, providing advice on experimental design and data interpretation. **H.M.R.R:** Shaping content, providing advice on experimental design and data interpretation. **M.R:** Conceptualisation, supervision, project administration, writing–review and editing. **G.S.H:** Conceptualisation, supervision, project administration, writing–review and editing.

## ACKNOWLEDGEMENTS

This work was primarily funded by the CRUK RadNet Oxford Centre (Cancer Research UK C6078/A28736). **P.T:** was supported by a Medical Research Council studentship (W006731/1)/(2744412). **G.S.H:** was also supported by a permanent endowment from the Howat Foundation.

## ETHICS DECLARATIONS

Competing interests

Al.Ci, M.L.B, Au.Ce, V.G, K.K, J.B.M, Su.Me, J.N, E.R, M.R.L, A.T, R.A.H, **G.C.M.S, H.M.R.R, M.R:** are all employees and shareholders of Artios Pharma Ltd. **M.L.B, Sh.Mo, B.R.H, S.A.L:** are former employees and shareholders of Artios Pharma Ltd. **Su.Me:** is a shareholder of AstraZeneca PLC. **G.C.M.S:** is a shareholder of AstraZeneca PLC **H.M.R.R:** is a shareholder of Mission Therapeutics. **A.T:** has share options in Benevolent AI. **J.B.M:** is a shareholder of AstraZeneca PLC and Crescendo. **E.R, R.A.H, G.C.M.S, H.M.R.R, M.R:** all hold patents with Artios Pharma Ltd. **G.C.M.S:** also reports a patent with AstraZeneca PLC. **H.M.R.R:** also reports a patent with Mission Therapeutics. **J.B.M:** is a patent holder with Crescendo. **G.S.H:** reports grants from Cancer Research UK, as well as personal fees and non-financial support from Artios Pharma during the conduct of the study. **G.S.H:** also reports personal fees from Exscientia, grants and personal fees from Sitryx, and grants from STipe Therapeutics outside the submitted work. **G.S.H:** is also a shareholder of Artios Pharma Ltd. No disclosures were reported by the other authors.

All other authors declare no competing interests.

