## Supplementary Figures, Methods and Tables for "SHLD2 loss is a synthetic vulnerability to Polθ inhibition combined with radiotherapy"

This PDF file contains:

1. Supplementary Figures 1-5
2. Supplementary Methods
3. Supplementary Tables 1-4

**Supplementary Figure 1 (accompanying Figure 1).**

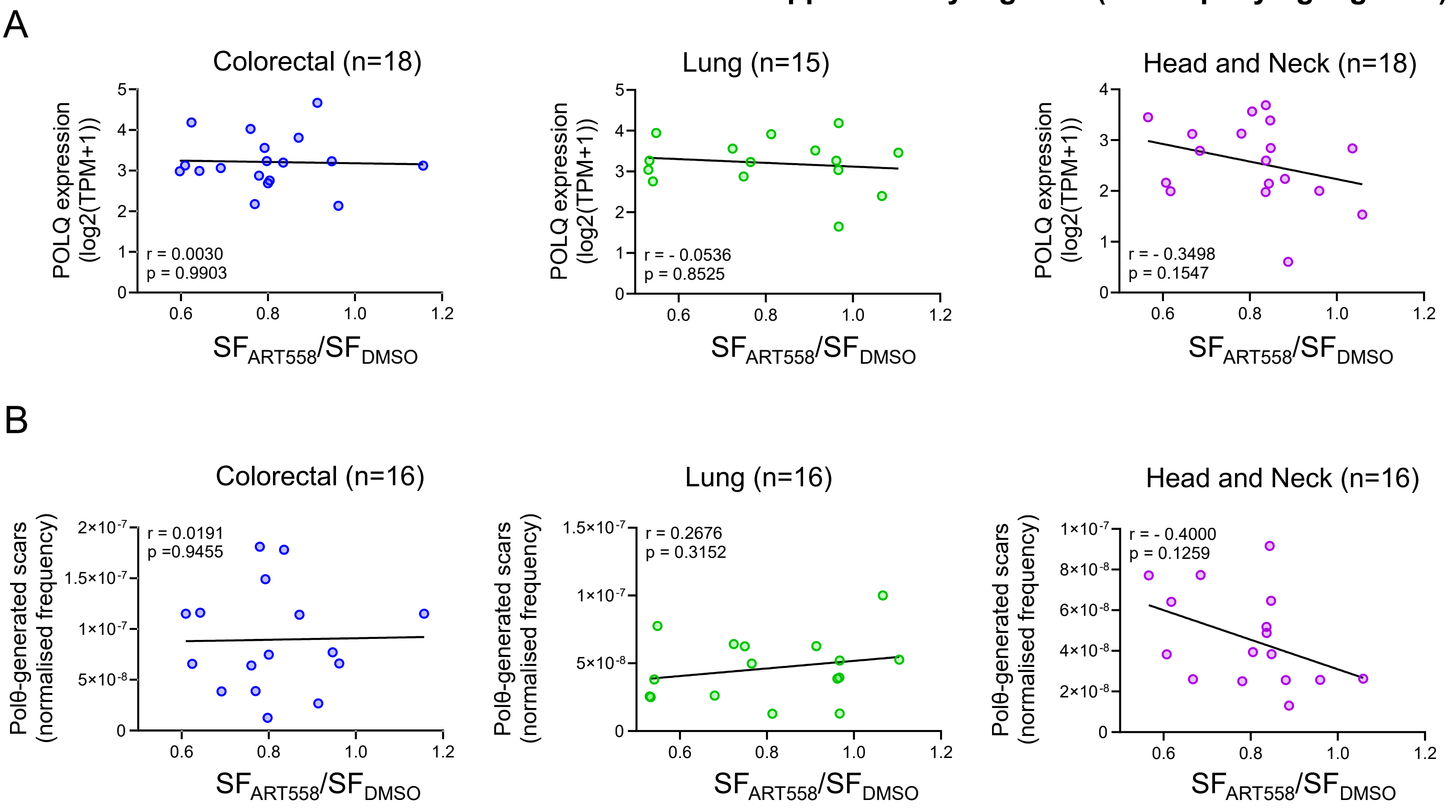


**Supplementary Figure 1 (accompanying Figure 1).**

(A) Correlation between *POLQ* expression and radiosensitisation by Polθi across tumour types.

(B) Correlation between the frequency of Polθ-generated scars and radiosensitisation by Polθi across tumour types.

Dots represent individual cell lines. Correlation coefficients (r) were calculated using the Spearman's method.

The black lines are regression lines.

**Supplementary Figure 2 (accompanying Figure 2).**

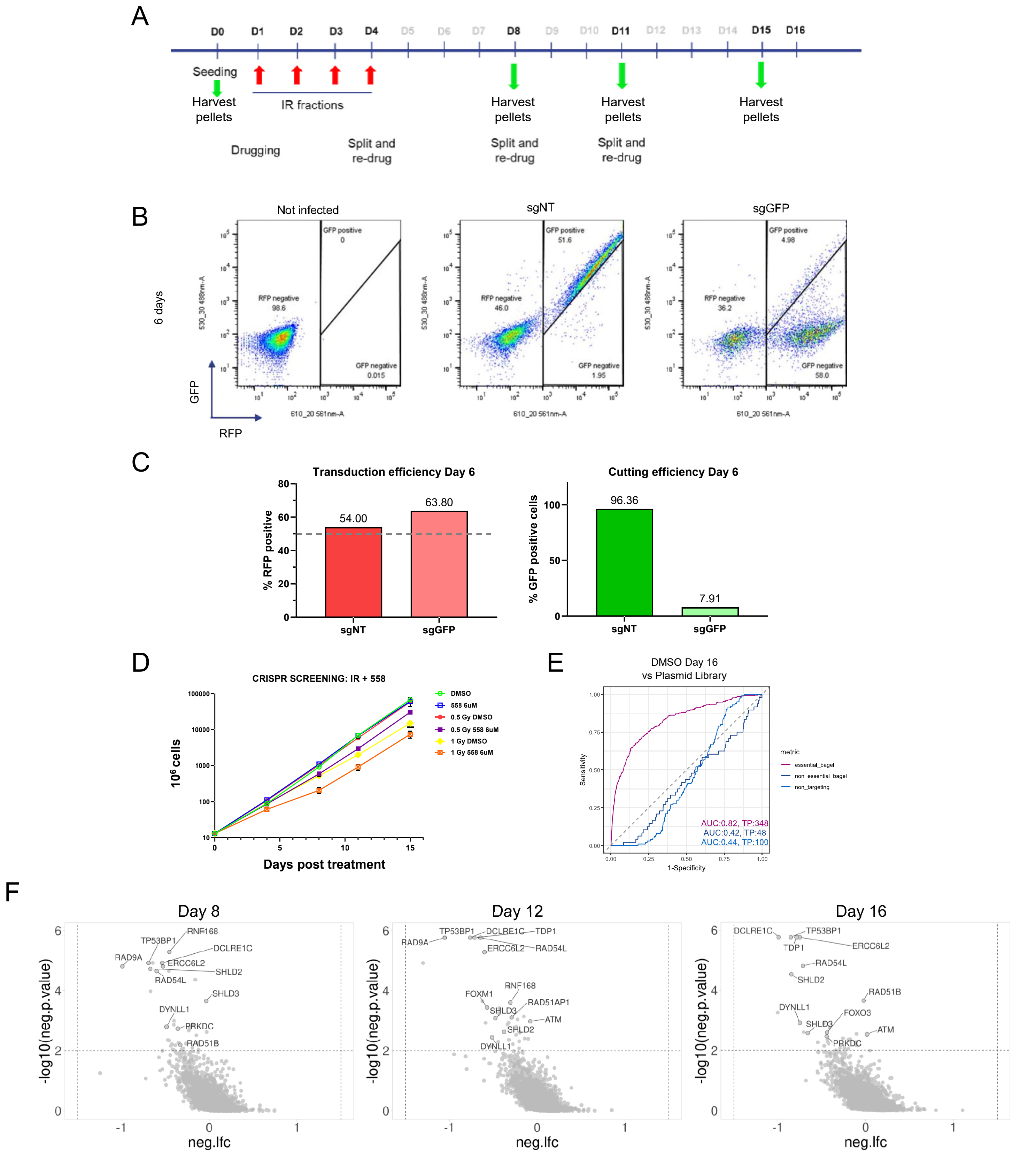

**Supplementary Figure 2 (accompanying Figure 2).**

(A) Experimental timeline of the CRISPR KO screen.

(B and C) Assessment of Cas9 cutting efficiency with CRISPRuTest$^{TM}$. Cas9-expressing DLD-1 cells were infected with a plasmid encoding for RFP and GFP, and either GFP-targeting sgRNA (sgGFP) or control sgRNA (sgNT). The higher the reduction in GFP signal analysed by FACS, the higher the cutting efficiency of Cas9.

(B) FACS plots showing Cas9 cutting efficiency. (C) Transduction efficiency and Cas9 cutting efficiency based on FACS plots shown in B.

(D) Growth curves of Cas9-expressing DLD-1 cells throughout the CRISPR screen.

(E) Receiver Operating Characteristic (ROC) curve showing sensitivity and specificity for essential and non-essential genes, and non-targeting sgRNAs for DMSO (day 16) vs plasmid library.

(F) Volcano plots highlighting the specific genes whose knockout synergised with combined Polθi (ART558) and RT (4 x 1 Gy) in at least two of the time points and/or RT schedules. neg.lfc: negative log fold change.

**Supplementary Figure 3 (accompanying Figure 3).**

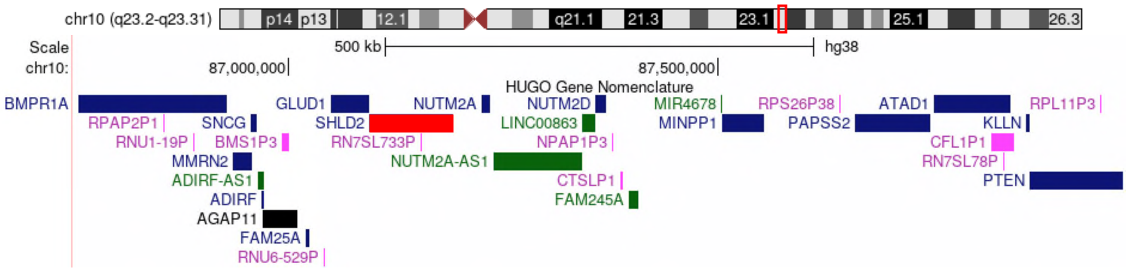


**Supplementary Figure 3 (accompanying Figure 3).**

UCSC Genome Browser visualisation of the genomic positions of *SHLD2* (red) and *PTEN* on chromosome 10 (GRCh38/hg38, Genome Reference Consortium).

**Supplementary Figure 4 (accompanying Figure 4).**

A

| Cell line | Allele 1 (bp) | Allele 2 (bp) | Predicted frameshift (%) | $R^2$ |
|---|---|---|---|---|
| CAL-51 clone E1 | -17 | -17 | 96 | 0.97 |
| CAL-51 clone E2 | -1 | -28 | 98 | 0.94 |
| DU145 clone A2 | -1 | -4 | 97 | 0.96 |
| 22Rv1 clone A6 | +1 | +8 | 96 | 0.94 |
| 22Rv1 clone C1 | +1 | -2 | 97 | 0.96 |
| 22Rv1 clone G6 | +1 | +1 | 97 | 0.98 |

B

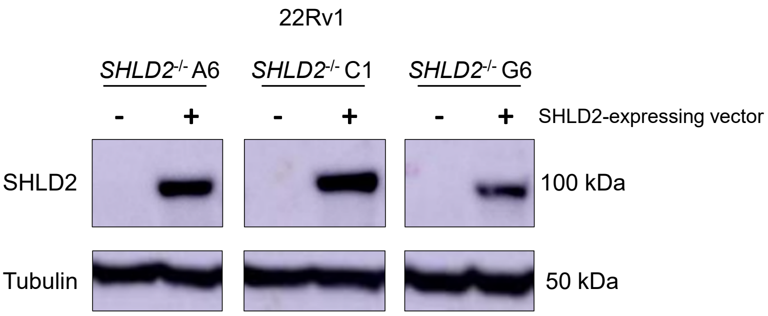


**Supplementary Figure 4 (accompanying Figure 4).**

(A) Confirmation of *SHLD2* KO status in the indicated cell lines. bp: base pair. (+) or (-) indicates an insertion or deletion, respectively.

(B) Western blot from lysates of *SHLD2*$^{-/-}$ 22Rv1 clones transduced with either a vector expressing *SHLD2* (+) or a GFP control vector (-).

| Mouse xenograft sample | Allele 1 (bp) | Allele 2 (bp) | Predicted frameshift (%) | $R^2$ |
|---|---|---|---|---|
| DU145 clone A2 | -1 | -4 | 96 | 0.97 |

**Supplementary Figure 5 (accompanying Figure 6).**

Confirmation of *SHLD2* KO status in the mouse xenografts. bp: base pair. (+) or (-) indicates an insertion or deletion respectively.

### Supplementary Methods

*SHLD2* knockout confirmation

The *SHLD2* PCR forward primer was CAAGGAGAGAGGACATGTTAGC (Sigma Aldrich SY230122662-075) and *SHLD2* PCR reverse primer was AGGGTTTACAGACTAATTTTTCCAG (Sigma Aldrich SY230122662-076). Genomic DNA was extracted using Lucigen QuickExtract™ DNA Extraction Solution (QE09050) and subjected to 65°C for 10 minutes, followed by 98°C for 5 minutes using a MasterCycler X50S. PCR was performed in duplicate samples using Q5 Hot Start High-Fidelity 2X Master Mix (New England Biolabs M0494S) in a Mastercycler X50 thermocycler. PCR cycling conditions were as follows: initial denaturation at 98°C for 30 seconds; 30 cycles of denaturation at 98°C for 10 secs; annealing at 63°C for 30 seconds; and extension at 72°C for 30 secs; final extension step at 72°C for 2 mins (see Supplementary Table 4). The PCR product was sequenced by Source Bioscience and the sequencing data was analysed using the ICE platform (Synthego, https://ice.editco.bio/#/) to obtain the indel frequency spectrum and determine the genetic make-up of the KO clones.

**Supplementary Table 1**. Cell lines, origin and culture media.

| Cell line ID | Source | Media + supplements |
|---|---|---|
| DU 145 | ATCC (HTB-81) | EMEM + 10% FBS |
| CAL-51 | DSMZ (ACC 302) | DMEM (high glucose) + 10% FBS |
| 22Rv1 | ATCC (CRL-2505) | RPMI 1640 + 10% FBS |
| Calu-6 | ATCC (HTB-56) | EMEM + 10% FBS |
| A549 | ATCC (CCL-185) | Ham's F-12K (Kaighn's) + 10% FBS |
| NCI-H2073 | ATCC (CRL-5918) | RPMI 1640 + 10% FBS |
| NCI-H1651 | ATCC (CRL-5884) | DMEM:F12 + 0.02 mg/mL insulin, 0.01 mg/mL transferrin, 25 nM sodium selenite, 50 nM Hydrocortisone, 1 ng/mL EGF, 0.01 mM ethanolamine, 0.01 mM phosphorylethanolamine, 100 pM triidothyronine, 0.5% BSA, 10 mM HEPES, 0.5 mM sodium pyruvate, 2 mM L-glutamine (final conc. 4.5 mM), 10% FBS |
| YD-15 | Horizon Discovery Inc. | RPMI + 10% FBS + 25 mM HEPES + 25 mM Sodium Bicarbonate |
| HT-115 | Horizon Discovery Inc. | DMEM + 15% FBS + 2 mM Glutamine |
| YD-8 | Horizon Discovery Inc. | RPMI + 10% FBS + 25 mM HEPES + 25 mM Sodium Bicarbonate |
| COLO-320 | Horizon Discovery Inc. | RPMI 1640 + 10% FBS |
| BICR 16 | Horizon Discovery Inc. | DMEM + 10% FBS + 0.4 µg/mL Hydrocortisone |
| HCT-15 | Horizon Discovery Inc. | RPMI 1640 + 10% FBS |
| DLD-1 | ATCC (CCL-221) | RPMI 1640 + 10% FBS |
| YD-10B | Horizon Discovery Inc. | RPMI + 10% FBS + 25 mM HEPES + 25 mM Sodium Bicarbonate |
| SK-LU-1 | ATCC (HTB-57) | EMEM + 10% FBS |
| CAL-33 | DSMZ (ACC447) | DMEM (high glucose) + 10% FBS |
| SW620 | Horizon Discovery Inc. | RPMI 1640 + 10% FBS |
| NCI-H1568 | ATCC (CRL-5876) | RPMI 1640 + 10% FBS |
| NCI-H1299 | ATCC (CRL-5803) | RPMI 1640 + 10% FBS |
| NCI-H747 | Horizon Discovery Inc. | RPMI 1640 + 10% FBS |
| SW900 | ATCC (HTB-59) | RPMI 1640 + 10% FBS |
| MDST8 | Horizon Discovery Inc. | DMEM + 10% FBS |
| RCM-1 | Horizon Discovery Inc. | RPMI:Ham’s F12 (1:1) + 10% FBS |

| | | |
|---|---|---|
| Detroit562 | Horizon Discovery Inc. | EMEM + 10% FBS |
| RKO | Horizon Discovery Inc. | EMEM + 10% FBS |
| HT29 | Horizon Discovery Inc. | McCoy’s 5A + 10% FBS |
| SW48 | Horizon Discovery Inc. | RPMI 1640 + 10% FBS |
| HCT-8 | Horizon Discovery Inc. | RPMI + 10% Horse Serum |
| SW480 | Horizon Discovery Inc. | RPMI 1640 + 10% FBS |
| PE/CA-PJ15 | Horizon Discovery Inc. | IMDM + 10% FBS |
| MORCPR | ECACC (Sigma-Merck) | RPMI 1640 + 10% FBS + 1 µg/mL cisplatin |
| KM12 | Horizon Discovery Inc. | RPMI 1640 + 10% FBS |
| SNU-1066 | Horizon Discovery Inc. | ATCC-formulated RPMI + 10% FBS + 25 mM HEPES + 25 mM Sodium Bicarbonate |
| FaDu | Horizon Discovery Inc. | EMEM + 10% FBS |
| PE-CA-PJ41-cl D2 | Horizon Discovery Inc. | IMDM + 10% FBS |
| YD-38 | Horizon Discovery Inc. | RPMI + 10% FBS + 25 mM HEPES + 25 mM Sodium Bicarbonate |
| HSC-3 | Horizon Discovery Inc. | EMEM + 10% FBS |
| A253 | ATCC (HTB-41) | McCoy's 5A Medium + 10% FBS |
| LS-411N | Horizon Discovery Inc. | RPMI 1640 + 10% FBS |
| HSC-4 | Horizon Discovery Inc. | EMEM + 10% FBS |
| BICR 56 | Horizon Discovery Inc. | DMEM + 10% FBS + 0.4 µg/mL Hydrocortisone |
| NCI-H1650 | ATCC (CRL-5883) | RPMI 1640 + 10% FBS |
| SW837 | Horizon Discovery Inc. | RPMI 1640 + 10% FBS |
| WiDr | Horizon Discovery Inc. | EMEM + 10% FBS |
| LS-123 | Horizon Discovery Inc. | EMEM + 10% FBS |
| SNU-1076 | Horizon Discovery Inc. | ATCC-formulated RPMI + 10% FBS + 25 mM HEPES + 25 mM Sodium Bicarbonate |
| NCI-H292 | ATCC (CRL-1848) | RPMI 1640 + 10% FBS |
| HT55 | Horizon Discovery Inc. | EMEM + 20% FBS |

| | | |
|---|---|---|
| HCC-15 | DSMZ (ACC496) | RPMI 1640 + 10% FBS |
| NCI-H1395 | ATCC (CRL-5868) | RPMI 1640 + 10% FBS |
| NCI-H1373 | ATCC (CRL-5866) | RPMI 1640 + 10% FBS |
| CAL-27 | Horizon Discovery Inc. | DMEM + 10% FBS |
| BICR 6 | Horizon Discovery Inc. | DMEM + 10% FBS + 0.4 µg/mL Hydrocortisone |
| NCI-H1869 | ATCC (CRL-5900) | DMEM:F12 + 0.02 mg/mL insulin, 0.01 mg/mL transferrin, 25 nM sodium selenite, 50 nM Hydrocortisone, 1 ng/mL EGF, 0.01 mM ethanolamine, 0.01 mM phosphorylethanolamine, 100 pM triidothyronine, 0.5% BSA, 10 mM HEPES, 0.5 mM sodium pyruvate, 2 mM L-glutamine (final conc. 4.5 mM), 10% FBS |
| NCI-H441 | ATCC (HTB-174) | RPMI 1640 + 10% FBS |
| LoVo | Horizon Discovery Inc. | Ham’s F12K + 10% FBS |

**Supplementary Table 2**. Reagents and concentrations used to produce CSK buffer for immunofluorescence microscopy.

| Reagent | Supplier | Reference | Final concentration |
|---|---|---|---|
| PIPES pH6.8 | Thermo Fisher | J61786.AE | 10 mM |
| NaCl | Sigma Aldrich | S9888-500G | 100 mM |
| Sucrose | Sigma Aldrich | S0389-500G | 300 mM |
| MgCl2 | Sigma Aldrich | 208337-100G | 1.5 mM |
| EDTA | Merck | 324506 | 5 mM |
| Protease inhibitor | Merck | 11873580001 | 1 tablet per 10 mL |
| Phosphatase inhibitor | Merck | 4906837001 | 1 tablet per 10 mL |
| Triton-X100 | Merck | X100-100 mL | 0.5% |

**Supplementary Table 3**. Antibodies used for immunofluorescence microscopy.

| Primary Antibodies | Species | Manufacturer; Product code | Dilution Factor | Blocking buffer |
|---|---|---|---|---|
| γH2AX | Mouse | Millipore; 05-636 | 1:2,000 | 0.5% BSA, 0.5% Triton |
| 53BP1 | Rabbit | Bethyl; A300-272A | 1:2,000 | 0.5% BSA, 0.5% Triton |
| pATM | Mouse | Millipore; 10h11.e12 | 1:5,000 | 0.5% BSA, 0.5% Triton |
| RAD51 | Rabbit | Millipore; ABE257 | 1:1,500 | 0.5% BSA, 0.5% Triton |
| **Fluorescent Secondary Antibodies** | **Species** | **Manufacturer; Product code** | **Dilution Factor** | **Blocking buffer** |
| DAPI | n/a | Sigma; D9542-5MG | 1:5000 | 0.5% BSA, 0.5% Triton |
| AlexaFluor 488 | Anti-Mouse | Thermo; A-11001 | 1:2,000 | 0.5% BSA, 0.5% Triton |
| AlexaFluor 568 | Anti-Rabbit | Thermo; A-21245 | 1:2,000 | 0.5% BSA, 0.5% Triton |

**Supplementary Table 4.** Thermocycler conditions for SHLD2 primer set PCR.

| Step | Temperature | Time |
|---|---|---|
| Initial Denaturation | 98°C | 30 seconds |
| 25-35 cycles | 98°C | 10 seconds |
| | 63°C | 30 seconds |
| | 72°C | 30 seconds |
| Final Extension | 72°C | 2 minutes |
| Hold | 4-10°C | |
